# Dysregulation of miR-103a-3p/AGFG1 axis links endocytic trafficking dysfunction to ER stress in Parkinson’s disease

**DOI:** 10.64898/2026.10.21.760838

**Authors:** Zhi-Wei Zhang, Sook-Yoong Chia, Lap Jack Wong, Zhihong Li, Haitao Tu, Eng-King Tan, Zhidong Zhou, Mian Cao, Li Zeng

**Author notes:** **Correspondence to:** Li Zeng, Neural Stem Cell Research Lab, Research Department, National Neuroscience Institute (NNI), 308433, Singapore, Singapore., Mian Cao, Neuroscience & Behavioral Disorders Program, Duke-NUS Medical School, 169857, Singapore, Singapore. These authors contributed equally to this work (Zhi-Wei Zhang, Sook-Yoong Chia).

## Abstract

**Background:** Parkinson’s disease is defined by progressive dopaminergic neurodegeneration and α-synuclein pathology. Endoplasmic reticulum (ER) stress and clathrin-mediated endocytosis dysfunction are implicated in disease pathogenesis, but the molecular mechanisms connecting these processes remain poorly understood. Here, we investigate a miR-103a-3p/AGFG1 axis as a regulator link between endocytic trafficking and ER stress in Parkinson’s disease.

**Methods:** MiR-103a-3p and AGFG1 were examined in plasma and postmortem substantia nigra from patients with Parkinson’s disease and controls. Gain-and loss-of-function studies, target validation and live-cell imaging were used to define the role of the miR-103a-3p/AGFG1 axis in ER stress and endocytic trafficking. The pathway was further investigated in SNCA A53T and LRRK2 G2019S hiPSC-derived neurons. Therapeutic targeting of AGFG1 was evaluated using antisense oligonucleotides (ASOs) *in vitro* and in a 6-hydroxydopamine mouse model.

**Results:** MiR-103a-3p was significantly reduced in Parkinson’s disease plasma and substantia nigra, whereas AGFG1 was increased in nigral tissue. MiR-103a-3p directly repressed AGFG1 and attenuated thapsigargin-induced ER stress, while miR-103a-3p inhibition or AGFG1 overexpression exacerbated stress signalling. AGFG1 knockdown reversed the effects of miR-103a-3p inhibition, establishing AGFG1 as a functional downstream effector. Mechanistically, AGFG1 associated and co-trafficked with clathrin-positive vesicles, whereas ER stress progressively disrupted AGFG1-clathrin vesicle dynamics, revealing an association between endocytic trafficking and stress signalling. In genetic Parkinson’s disease models, SNCA A53T and LRRK2 G2019S neurons showed mutation-dependent differences in AGFG1 expression and stress regulation. eIF2α phosphorylation increased in both SNCA A53T and LRRK2 G2019S neurons, whereas AGFG1 and GADD34 upregulation was selective to SNCA A53T neurons. Importantly, ASO-mediated AGFG1 suppression attenuated ER stress *in vitro* and preserved nigral dopaminergic neurons with improved motor performance *in vivo*.

**Conclusion:** Our findings identify the miR-103a-3p/AGFG1 axis as a previously unrecognised molecular link between endocytic trafficking dysfunction and ER stress in Parkinson’s disease, and support AGFG1 as a mechanistically potential therapeutic target.

## Introduction

Parkinson’s disease is a progressive neurodegenerative disorder, pathologically characterised by the accumulation of misfolded alpha-synuclein (-syn) in the Lewy bodies and loss of dopaminergic (DA) neurons in the substantia nigra (SN)^1^. Clinically, individuals with Parkinson’s disease manifests primarily as motor symptoms, including uncontrolled tremors, muscular rigidity, bradykinesia and postural instability, and is frequently accompanied by non-motor symptoms including cognitive impairment^2^. Current therapeutic approaches, such as dopamine replacement therapy and deep brain stimulation, provide symptomatic relief^3,4^ but neither halt neurodegeneration nor modify the disease progression. Moreover, long-term treatment may be associated with adverse effects^5^. Thus, there remains a critical unmet need to identify molecular mechanisms underlying dopaminergic neurodegeneration and to develop neuroprotective therapeutic strategies.

Accumulating evidence implicates endoplasmic reticulum (ER) stress as a critical contributor to dopaminergic neurodegeneration in Parkinson’s disease. Misfolded-syn aggregates have been detected within the ER lumen of patient tissues and experimental models, disrupting ER homeostasis and protein quality control mechanism^6^. Consistent with these observations, elevated expression of ER stress markers and unfolded protein response (UPR) has been reported in Parkinson’s disease, suggesting sustained ER dysfunction in disease pathogenesis^7–10^. ER stress arises when misfolded or unfolded proteins accumulate in the ER, triggering activation of the UPR through three primary stress transducers: protein kinase RNA-like ER kinase (PERK), inositol-requiring protein-1 (IRE1), and activating transcription factor-6 (ATF6)^11^. Although transient UPR activation is initially neuroprotective by restoring proteostais^12^, prolonged or excessive ER stress can shift UPR signaling toward pro-apoptotic pathways including induction ofs downstream chaperones C/EBP homolous protein (CHOP) and activation of caspases, ultimately contributing to cell death^13^. Understanding the molecular mechanisms that regulate ER stress responses may therefore reveal therapeutic targets for protecting dopaminergic neurons in Parkinson’s disease.

MicroRNAs (miRNAs) are small non-coding RNAs that post-transcriptionally regulate gene expression and play important roles in neuronal homeostasis and neurodegenerative processess. Through regulation of their target genes, miRNAs influence diverse cellular processes including neuronal differentiation, synaptic plasticity, apoptosis, and stress responses^14,15^. Their present in circulating biofluids, including plasma, supporting their potential utility as minimally invasive biomarkers for Parkinson’s disease^16^. Emerging evidence suggests bidirectional interactions between miRNA-mediated gene regulation and endoplasmic reticulum (ER) stress pathways, whereby miRNAs can modulate unfolded protein response (UPR) signalling and are themselves regulated by ER stress^17^. Previous studies have demonstrated the involvement of miRNA-mediated pathways in ER stress regulation and dopaminergic neuronal injury in Parkison’s disease^18–20^. However, the potential involvement of miRNA-mediated regulation in connecting ER stress with endocytic trafficking and dopaminergic neurodegeneration remain incompletely understood.

Among miRNAs potentially involved in neurodegeneration, miR-103a-3p is abundantly expressed in the human brain^21^. Previous miRNA profiling of postmortem brain tissues from patients with Parkinson’s disease revealed reduced precursor miR-103 expression in both cerebral cortex and mibrain compared with controls^22^. This observation provides the initial rationale for investigating the functional significance of mature miR-103a-3p in Parkins’s disease pathogenesis. Beyond Parkinson’s disease, miR-103a-3p has been implicated in diverse pathological processes, including inflammatory reponses^24,25^, renal injury^26^, Deep Vein Thrombosis^27^, cancer^28–30^ and Alzheimer’s disease^31^. Despite these associations, the functional significance and molecular mechanisms of of miR-103a-3p in Parkinson’s disease remain largely unexplored. In particular, whether miR-103a-3p contributes to dopaminergic neuron vulnerability through ER stress is unknown.

Motivated by the reported reduction of pre-miR-103 in the brain of Parkinson’s disease, we investigated the expression, functional significance and molecular mechanisms of miR-103a-3p in Parkinson’s disease. We showed that miR-103a-3p is consistently downregulated in both the plasma and the SN of Parkinson’s disease. Using dopaminergic SH-SH5Y neurons as an *in vitro* Parkinson’s disease model, we demonstrated that miR-103a-3p suppresses sustained ER stress induced by thapsigargin (TG). Mechanistically, we identify Arf-GAP with FG repeats protein 1 (AGFG1), a regulator of clathrin-mediated endocytosis (CME)^32^, as a direct downstream target of miR-103a-3p. We further reveal that dysregulation of the miR-103a-3p/AGFG1 axis impaired endocytic dynamics and exacerbated ER stress signaling. Importantly, pharmacological suppression of AGFG1 using antisense oligonucleotides (ASOs) attenuated ER stress, preserved nigral dopaminergic neurons and improved motor function in a Parkinson’s disease mouse model. Together, our findings identify a previously unrecognized miR-103a-3p/AGFG1 regulatory axis associated with altered endocytic trafficking and ER stress in Parkinson’s disease. These findings provide mechanistic insights into dopaminergic neurodegeneration and highlight AGFG1 as a potential therapeutic target..

## Materials and methods

### Study participants and plasma collection

A total of 15 patients with Parkinsons’s disease and 16 age-and sex-matched healthy controls (HCs) were included in this case-control study, with a mean age ranging from 62-67 years across the groups (**Table 1**). Participants were recruited from Singpore General Hospital, National Neuroscience Institute, affliated SingHealth clinics, as well as inpatient wards, community outreach programmes, the Clinical Trials and Research Centre (CTRC) healthy volunteer registry, and participants referrals. Parkinson’s disease Diagnosis was diagnosed according to National Institute of Neurological Disorders and Stroke (NINDS) diagnostic criteria^33^. Our study was approved by the SingHealth Centralized Institutional Review Board (Ref 2019/2330), and written informed consent was provided by all participants. All procedures were conducted in accordance with the approved guidelines and regulations. Blood was collected in EDTA tubes and centrifuged at 1,500 x g for 10 min at 4. Plasma was aliquoted and stored at −80 until analysis.

**Table 1.** Demographic of patients with Parkinson’s disease and healthy controls. Values are given as n (%) or mean (SD). Differences between groups were assessed using Fisher’s exact test (sex) and Mann-Whitney *U-*test (age and age at onset).

|  | Healthy control | Parkinson's disease | *p*-value |
| --- | --- | --- | --- |
| No of subjects | 15 | 16 |  |
| Age, years | 62.33 (8.41) | 67.00 (11.80) | 0.1973 |
| Sex |  |  |  |
| Male | 12 (75%) | 10 (66.67%) |  |
| Female | 4 (25%) | 5 (33.33%) | 0.7043 |
| Age at onset, years | - | 65.53 (11.97) |  |

### Human human substantia nigra tissues

Postmortem substantia nigra (SN) tissues from 8 individuals with Parkinson’s disease and 8 age-matched HCs were obtained from the Parkinson’s UK Brain Bank, Division of Neuroscience, Imperial College London, as previously described^34,35^. Tissue samples were analysed by Western blot and qPCR. The demographic characteristics and neuropathological diagnoses of the cases are summarised in **Supplementary Table S1** and **Table S2**, respectively. All tissues were obtained via a prospective donor scheme with fully written consent, and their collection was approved by the Human Tissue Authority (Approval Number 18/WA/0238). The study was also approved by the SingHealth Institutional Review Board Committee and conducted in accordance with the approved guidelines and regulations.

### Primary cortical neuron culture and immunofluorescence

Cultures of cortical neurons were prepared from P0 to P2 neonatal mouse brains by previously described methods^36^ and used at 19 days *in vitro* (DIV). Cortical tissue was dissected out, placed in ice-cold Hanks’ Balanced Salt Solution (HBSS), and diced into small pieces of less than 1 mm^3^ per piece. Tissue was then digested for 30 min in an activated enzyme solution containing papain (20 U/ml) and DNase (20 µg/ml) at 37°C, followed by gentle trituration. The cell suspension is centrifuged at 300 g for 5 min. The cell pellet is resuspended and plated onto poly-d-lysine-coated coverslips to a density of 60,000 cells/cm^2^. 2-3 h after plating, the medium was exchanged to Neurobasal/B27 serum-free medium, and cells were maintained at 37°C in a 5% CO_2_ humidified incubator. Cells were fixed with 4% paraformaldehyde (PFA) and 4% sucrose in 1x PBS for 20 min, followed by 10 min incubation with 50 mM NH_4_Cl, then blocked and permeabilised with 5% bovine serum albumin (BSA), 1x phosphate-buffered saline (PBS) and 0.1% Triton X-100. Primary and secondary antibody incubations were subsequently performed in the same buffer. Alexa 488, 594 and 647 conjugated secondary antibodies were purchased from Invitrogen. After washing, samples were mounted on slides with Fluoromount-G (Invitrogen). Samples were observed and imaged on a spinning disk system (Gataca Systems) based on an inverted microscope (Nikon Ti2-E; Nikon) equipped with a confocal spinning head (CSU-W, Yokogawa) and a Plan-Apo 60x oil TIRF objective. The same exposure time and laser intensity were used when imaging the same marker for all genotypes.

### Antisense Oligonucleotides (ASOs)

AGFG1-targeting ASOs were purchased from Integrated DNA Technologies (IDT) and dissolved in nuclease-free water. Two functional ASOs regions were identified. ASO1: CGACTTTGGCTTGTTCTGGC. ASO2: ACGACCTACAACTGGGGACT.

### Cell culture, transfection and live-cell imaging

SH-SY5Y and HEK293 cells (ATCC) was cultured in DMEM containing Fetal Bovine Serum (Hyclone), 1% penicillin-streptomysin (Gibco), and 1% L-GlutaMAX (Gibco) at 37°C in 5% CO_2_ humidified incubator. Transfection of plasmids or miRNAs were performed using Lipofectamine LTX (Invitrogen) according to the manufacturer’s instructions. The miRNAs and plasmid constructs used as described in **Supplemental method**.

For live-cell imaging, SH-SY5Y cells were transfected with GFP-AGFG1 and RFP-AP2 and imaged 48 h later using a spinning disk system (Gataca Systems) based on an inverted microscope (Nikon Ti2-E; Nikon) equipped with a confocal spinning head (CSU-W, Yokogawa), a Plan-Apo 60x oil TIRF objective, a Tokai Hit INU-NI-F1 temperature, humidity, and CO_2_ chamber. Time-lapse images were acquired at 5 seconds intervals with 400 ms exposures. Image anlaysis were performed using MetaMorph (Universal Imaging) and Image J software (NIH).

Stable AGFG1 overexpression and knockdown SH-SY5Y cells were generated by transfection with pCAG-BSD-AGFG1::HAP2A::EGFP for 48 h or pGreenPuro-shRNA-AGFG1, respectively, followed by 2 μM puromycin selection.

### Thapsigargin treatment

Thapsigargin (TG, Sigma) was dissolved in DMSO (Sigma) to serve as the vehicle. SH-SY5Y cells were seeded in 6-well plates (100,000 cells/well), or onto poly-L-lysine-coated coverslips (20,000 cells/coverslip) and slowed to incubate for 24 h. Following this, the cells were transfected with miRNA or plasmids using Lipofectamine LTX and incubated for additional 48 h, concurrently treated with 1 μM TG for varying duration of 2-24 h prior to downstream analyses.

### RNA extraction and qPCR

Total RNAs including miRNAs were extracted from plasma using miRNeasy Serum/Plasma Kit (Qiagen) and from SN tissues or transfected cells using miRNeasy Mini Kit (Qiagen). miRNA expressions were validated by using Taqman MicroRNA Assays (Applied Biosystem) following the procedure described previously^37^.

For mRNA analysis, reverse transcription was performed using iScript^TM^ cDNA Synthesis kit (Bio-rad), followed by qPCR using All-in-One^TM^ qPCR Mix (Genecopoeia). β-actin was used as a reference gene. Relative expression of the genes were calculated using the 2^−ΔΔCT^ method relative to values of β-actin.

### Western blotting

Proteins were extracted using RIPA lysis buffer supplemented with protease and phosphatase inhibitor cocktail (MedChemExpress). Protein concentrations were measured using the RC-DC protein assay kit II (Bio-Rad). Equal amounts of protein lysates (20 μg) were separated through SDS-PAGE and then transferred to a polyvinylidene fluoride membrane (Millipore). Membranes were blocked for 1 h at room temperature using 5% milk in tris buffered saline with tween 20 (TBST) and incubated overnight with a 1:1000 dilution of primary antibodies (**Supplementary Table**) at 4°C. After incubation with horseradish peroxidase-conjugated secondary antibody (1:3000, GE Healthcare), signals were detected using Pierce ECL Western Blotting Substrate and SuperSignal West Femto Maximum Sensitivity Substrate (Thermofisher Scientific). Band intensity was quantified using ImageJ and normalised β-actin (Santa Cruz).

### TUNEL assays

Apoptotic cell death was assessed using In Situ Cell Death Detection Kit (Roche) according to the manufacturer’s instructions. Images were captured using confocal microscopy (FV3000 Olympus).

### Luciferase reporter assay

To elucidate the putative complementary sequences of miR-103a-3p identified in the 3′ untranslated region (3’ UTR) of AGFG1 mRNA, wild-type (AGFG1-WT) and mutant (AGFG1-MT) fragments were inserted into luciferase reporter constructs. HEK293 cells were co-transfected with miRNA and luciferase plasmids using the Lipofectamine LTX. Luciferase activities were measured using the Dual Luciferase Reporter Assay Kit (Promega).

### Human iPSC-derived neuron differentiation

Human induced pluripotent stem cell (hiPSC) lines harbouring homozygous SNCA A53T (KOLF2.1J; JIPSC001130) and LRRK2 G2019S (KOLF2.1J; JIPSC003262), together with their shared isogenic controls (KOLF2.1J; JIPSC00100), were were differentiated into pan-neuronal cells using established protocol^38–40^. Detailed differentiation procedures are provided in the **Supplementary method**.

### 6-OHDA lesion–induced Parkinson’s disease mouse model

Two-month-old male C57BL/6 mice were anesthetised with a mixture of ketamine (150 mg/kg; Ceva Animal Health Pty Ltd.) and xylazine hydrochloride (10 mg/kg; Troy Laboratories Pty Ltd.) and secured in a stereotaxic frame. Mice were randomly assign to be control or treatment groups. 6-hydroxydopamine (6-OHDA, 4 μg in 0.9% NaCl containing ascorbic acid of 0.2 mg/ml) was injected into the left striatum at the coordinates: anterior-posterior = 0.5 mm and medial-lateral = 1.8 mm relative to bregma and 3 mm below skull at the rate of 0.5 μl/min. 6-OHDA (4 μg) was stereotaxically injected into left striatum of two-month-old C57 (day 0). AGFG1-ASO2 (2 µg) was then administered into the left SN on day eight via stereotaxically injection.

All procedures were approved by the Institutional Animal Care and Use Committee (IACUC) at the Lee Kong Chian School of Medicine, Nanyang Technological University, Singapore. The studies adhered to the National Advisory Committee for Laboratory Animal Research (NACLAR) Guidelines. Mice were maintained in a pathogen-free facility and exposed to a 12-h light/dark cycle, with food and water available ad libitum.

## Statistics Analysis

Data was analysed using GraphPad Prism 10. Results are presented as means ± standard deviation unless otherwise specified; human samples data are presented as means ± standard error of the mean (SEM). Statistical comparison between two groups were analysed using two-tailed Student’s *t*-test. Multiple group comparisons were analysed by one-way or two-way ANOVA followed by Tukey’s post hoc test. A *p* value < 0.05 was considered statistically significant.

## Results

### MiR-103a-3p is reduced in Parkinson’s disease and directly represses AGFG1 through 3’UTR binding

MiR-103a-3p is abundantly expressed in the human brain and dyregulated in Parkinson’s disease. Despite its involvement in diverse pathological processes, its role in Parkinson’s disease, particularly in ER stress and dopaminergic neuron susceptibility remain unclear. We first examined miR-103a-3p expression in plasma and substantia nigra (SN) from patients with Parkinson’s disease and healthy controls (HC). qPCR analysis revealed a significant reduction in miR-103a-3p levels in the plasma of Parkinson’s disease patients compared with HC (89.60% decrease, *p* < 0.0001; **Fig. 1A**). Consistently, miR-103a-3p levels were significantly reduced in the SN of Parkinson’s disease (67.24% decrease, *p* = 0.0365; **Fig. 1B**).

**Figure 1.**
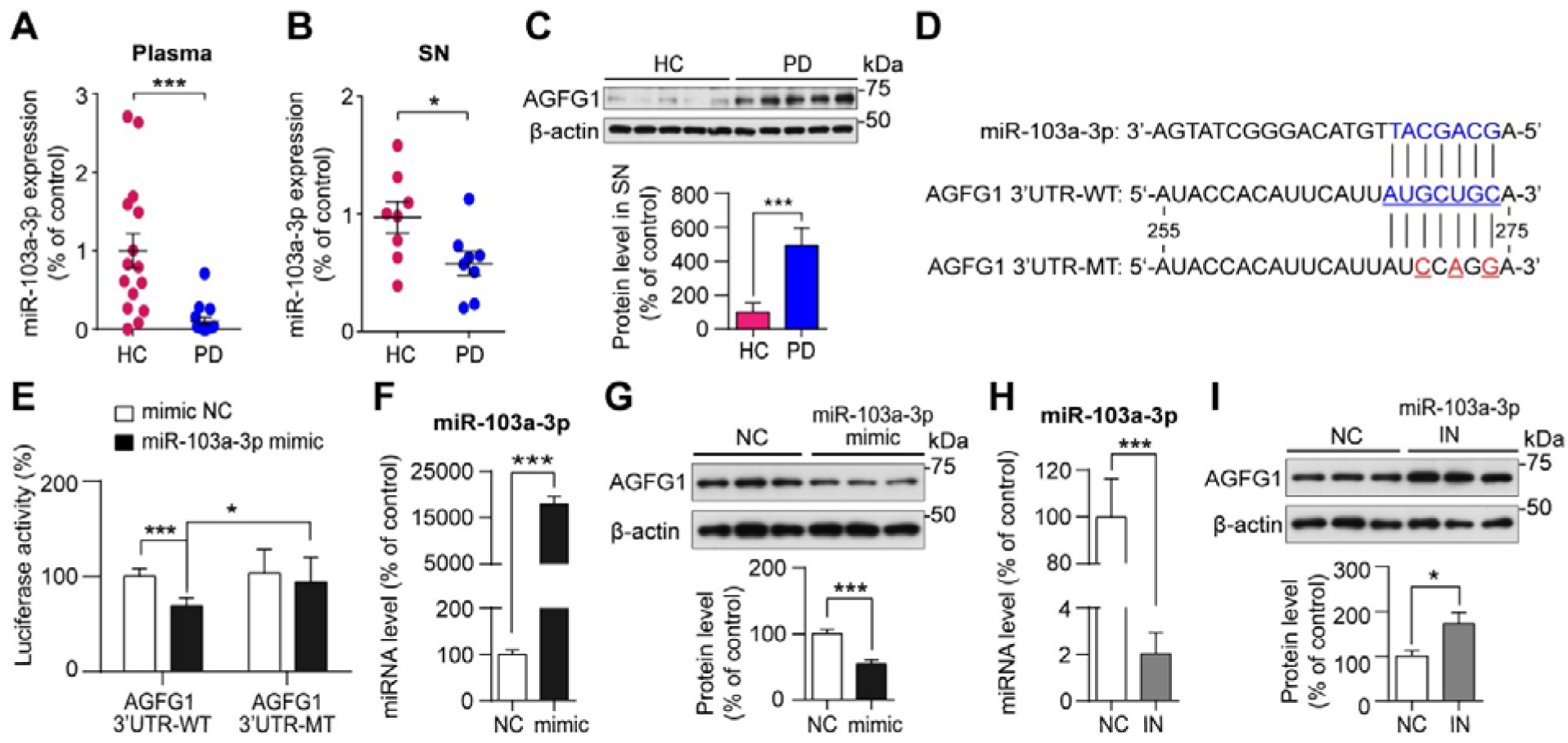
MiR-103a-3p is reduced in Parkinson’s disease and directly represses AGFG1 through 3’UTR binding. (**A**) miR-103a-3p expression are significantly reduced in plasma from patients with Parkinson’s disease (*n* = 16) compared with healthy controls (HC, *n* = 15). (**B**) MiR-103a-3p expression is similarly decreased in the SN of Parkinson’s disease subjects ( *n*= 8). Data are presented as mean ± SEM. (**C**) Representative Western blot and quantification of AGFG1 protein levels in the SN from Parkinson’s disease and HC (*n* = 8 per group). (**D**) Sequence analysis of human miR-103a-3p and the AGFG1 mRNA 3’UTR. The predicted miR-103a-3p binding site is highlighted in blue, and mutated nucleotides are shown in red. (**E**) Relative luciferase activity in HEK293T cells co-transfected with pmirGLO-WT-AGFG1-3’UTR wild-type (WT) or mutant (MT) constructs and either a control mimic or miR-103a-3p mimic (*n* = 6 per group). (**F-I**) SH-SY5Y cells were transfected with a miR-103a-3p mimic or inhibitor for 48 h. miR-103a-3p expression and representative Western blot of AGFG1 following (**F-G**) miR-103a-3p overexpression and (**H-I**) miR-103a-3p inhibition are shown (*n* = 3 per group). Data are presented as mean ± SD, \**p* < 0.05, \*\*\**p* < 0.001 by Student’s *t*-test.

To identify potential downstream targets of miR-103a-3p, we performed *in silico* target prediction miRWalk3.0, TargetScan, miRDB and miRTarBase^37,41^. Among the predicted targets, AGFG1 was priotised AGFG1 based on its high prediction ranking, two conserved 8-mer binding sites identified by TargetScan, a miRDB target score of 99 and previously reported experimentally validated miRNA-target interaction from CLASH and next generation sequencing (NGS) datasets curated in miRTarBase. Furthermore, AGFG1 has been implicated in CME^32^, a membrane-trafficking pathway is increasing reported to be associated with Parkinson’s disease^42,43^, whereas its potential involvement in miR-103a-3p-mediated ER stress regulation in Parkinson’s disease pathogenesis remains unexplored. Given the reduced miR-103a-3p expression observed in Parkinson’s disease, we next examined whether AGFG1 expression was correspondingly altered. AGFG1 protein levels were significantly increased in the SN of Parkinson’s disease patients (**Fig. 1C**). In contrast, AGFG1 mRNA levels remained unchanged in both plasma and SN samples (**Supplementary Fig. S1A-B**), indicating the increased AGFG1 protein expression may involve post-transcriptional regulation.

To experimentally validate the predicted miR-103a-3p binding sites within the AGFG1 3’UTR, we performed luciferase reporter assays in HEK293 cells co-transfected with miR-103a-3p and reporter constructs containing either the wild-type (WT) or mutant (MT) AGFG1 3’UTR in which the predicted miR-103a-3p binding site was disrupted (**Fig. 1D**). MiR-103a-3p overexpression significantly reduced luciferase activity in cells expressing the WT construct, whereas mutation of the predicted binding site abolished this effect (**Fig. 1E**). These results confirm that miR-103a-3p directly binds to the 3’UTR of AGFG1 to inhibit its expression.

We next examined whether modulation of miR-103a-3p altered endogenous AGFG1 expression in SH-SY5Y cells. Transfection efficiency of miR-103a-3p mimics or inhibitors was confirmed by qPCR (**Supplementary Fig. S1C**). Results showed that miR-103a-3p mimics significantly reduced AGFG1 protein levels (**Fig. 1F-G**) and AGFG1 mRNA levels (**Supplementary Fig. S1D**). Conversely, miR-103a-3p inhibition increased AGFG1 protein levels **(Fig 1H-I**) without altering AGFG1 mRNA levels (**Supplementary Fig. S1E**), consistent with observations in SN of Parkinson’s disease (**Fig. 1C**). These findings demonstrate that miR-103a-3p directly targets AGFG1 and negatively regulates its expression in in Parkinson’s disease.

### MiR-103a-3p suppresses thapsigargin-induced ER stress

To determine whether miR-103a-3p modulates the ER stress response, SH-SY5Y cells transfected with miR-103a-3p mimics or inhibitors or their respective negative controls were exposed to TG for 0, 2, 6, 12 and 24 h to induce ER stress. In NC-transfected cells, TG induced a transient increase in phosphorylated eukaryotic translation initiation factor 2α (P-eIF2α), whereas the expression of CHOP and (Growth Arrest and DNA-Damage-inducible protein 34) GADD34 progressively increased with prolonged TG exposure (**Fig. 2A-H**).

**Figure 2.**
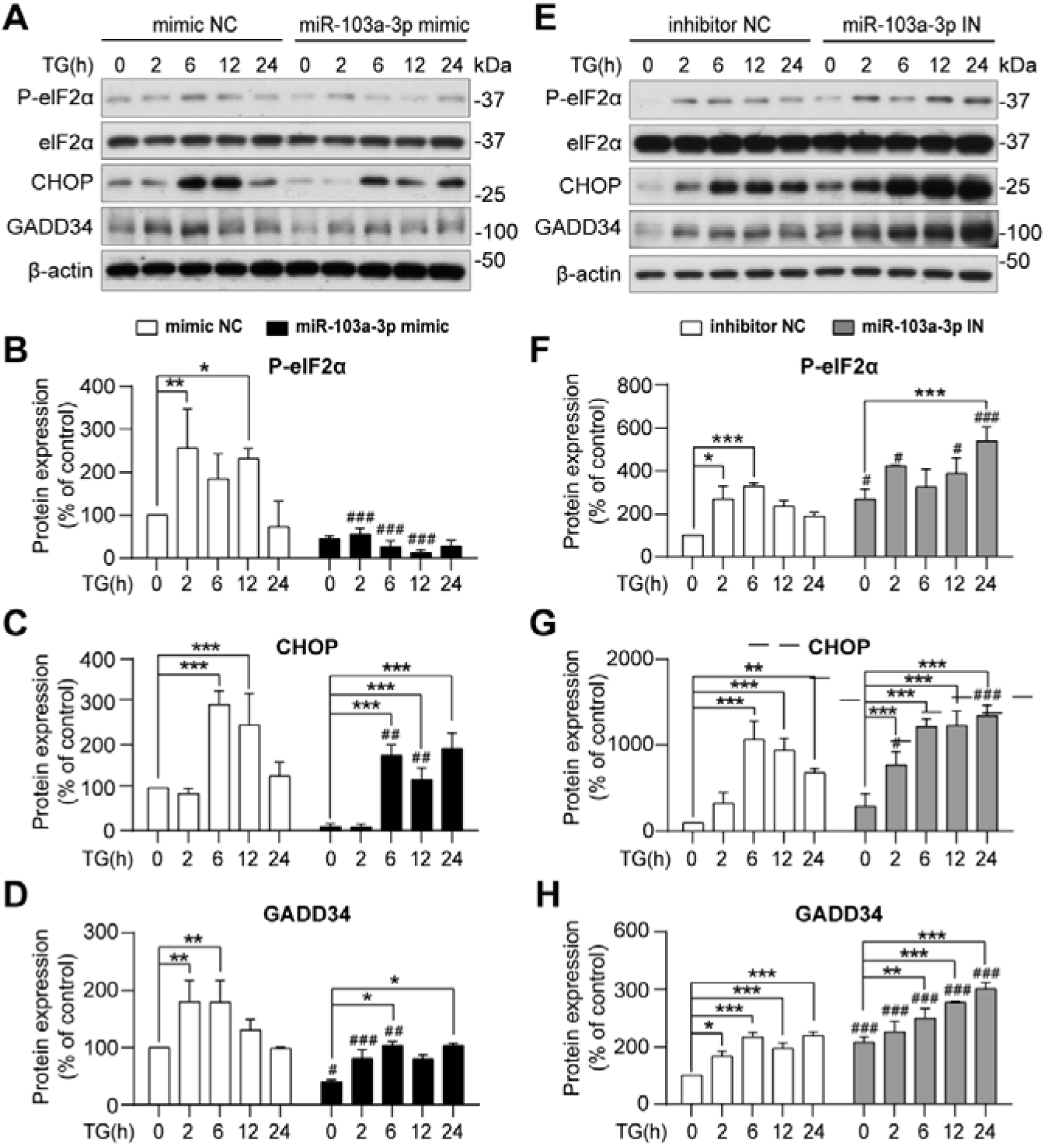
MiR-103a-3p suppresses thapsigargin-induced ER stress. (**A-D**) SH-SY5Y cells were transfected with miR-103a-3p mimic and control for 48 h, followed by treatment with 1 μM TG for 2, 6, 12 and 24 h. (**A**) Representative Western blot and quantitation of (**B**) P-eIF2α/eIF2α, a marker of unfolded protein response (UPR) activation, and downstream pro-apoptosis markers (**C**) CHOP and (**D**) GADD34. (**E-H**) SH-SY5Y cells transfected with miR-103a-3p inhibitor or control were treated with 1 μM TG for 2, 6, 12 and 24 h. (**E**) Representative Western blot and quantification of (**F**) P-eIF2α/eIF2α, (**G**) CHOP and (**H**) GADD34. Data are presented as mean ± SD (*n* = 3 per group). \**p* < 0.05, \*\**p* < 0.01, and \*\*\**p* < 0.001 relative to 0 h TG; ^#^*p* < 0.05, ^##^*p* < 0.01, and ^###^*p* < 0.001 relative to corresponding same timepoint of TG treatment. Statistical analysis was performed by two-way ANOVA with Tukey’s post hoc test.

MiR-103a-3p mimics attenuated TG-induced upregulation of P-eIF2α, CHOP and GADD34 compared with the corresponding NC (**Fig. 2A-D**). Conversely, miR-103a-3p inhibition enhanced the ER stress signaling in a time-dependent manner relative to the inhibitor NC (**Fig. 2E-H**). These results indicate that miR-103a-3p negatively regulates sustained ER stress signaling.

### AGFG1 exacerbates ER stress signaling

Having established AGFG1 as a direct target of miR-103a-3p (**Fig. 1**) and that miR-103a-3p modulates ER stress response (**Fig. 2**), we next examined whether AGFG1 contributes to ER stress regulation. SH-SY5Y cells with stable AGFG1 overexpression or knockdown were exposed to TG, and the temporal expression of ER stress markers was assessed. In control cells, TG treatment increased P-eIF2α, CHOP, and GADD34 expression, confirming ER stress induction (**Fig 3A-E**). AGFG1further increased P-eIF2α expression during the early phage of TG exposure (2 h) and enhanced CHOP and GADD34 expression following prolonged exposure (24 h). Conversely, AGFG1 knockdown significantly reduced the expression of ER stress marker during prolonged TG treatment (**Fig 3F-J**). Together, these results demonstrate that AGFG1 enhances both early UPR activation and downstream pro-apoptotic ER stress signaling.

**Figure 3.**
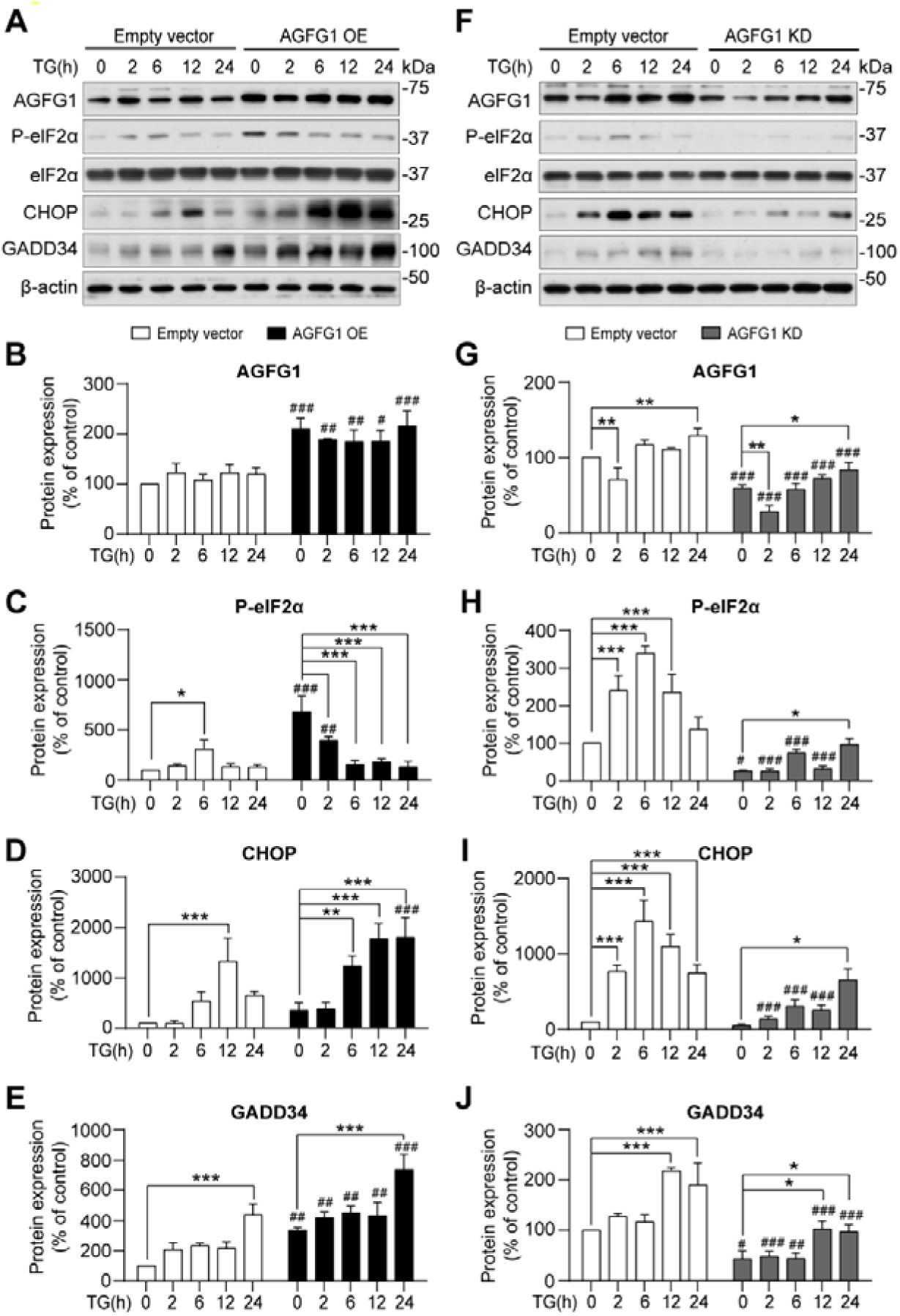
AGFG1 exacerbates ER stress signaling. (**A-E**) SH-SY5Y cells were transfected with *pCAG-BSD-AGFG1::HAP2A::EGFP* for 48 h in addition of 2 μM puromycin to establish the AGFG1 overexpression, and then treated with 1 μM TG for 2, 6, 12 and 24 h. (**A**) Representative Western blot and quantitation of (**B**) AGFG1, (**C**) P-eIF2α/eIF2α, (**D**) CHOP and (**E**) GADD34. (**F-J**) SH-SY5Y cells were transfected with *pGreenPuro-shRNA-AGFG1* in addition of 2 μM puromycin to establish the AGFG1 knockdown stable cell line and then treated with 1 μM TG for 2, 6, 12 and 24 h. (**F**) Representative Western blot and quantification of (**G**) AGFG1, (**H**) P-eIF2α/eIF2α, (**I**) CHOP and (**J**) GADD34. Data are presented as mean ± SD (*n* = 3 per group). \**p* < 0.05, \*\**p* < 0.01, and \*\*\**p* < 0.001 relative to 0 h TG; ^#^*p* < 0.05, ^##^*p* < 0.01, and ^###^*p* < 0.001 relative to corresponding same timepoint of TG treatment. Statistical analysis was performed by two-way ANOVA with Tukey’s post hoc test.

### AGFG1 knockdown rescues ER stress induced by miR-103-3p inhibition

To determine whether AGFG1 mediates the effects of miR-103a-3p on ER stress, control and AGFG1-knockdown SH-SY5Y cells were transfected with a miR-103a-3p inhibitor or inhibitor NC, followed by TG treatment for 24 h. In vector control cells, miR-103a-3p inhibition increased AGFG1expression, whereas this increase was abolished in AGFG1 knockdown cells (**Fig. 4A-B**). Moreover, miR-103a-3p inhibitionalso increased P-eIF2α, CHOP, and GADD34 expression following TG treatment. Important, these effects were significantly attenuated by AGFG1 knockdown (**Fig. 4A, C-E**). These findings suggest that AGFG1 contributes to the enhancement of ER stress signaling induced by miR-103-3p inhibition and support AGFG1 as a downstream efffector of miR-103a-3p-mediated ER stress regulation.

**Figure 4.**
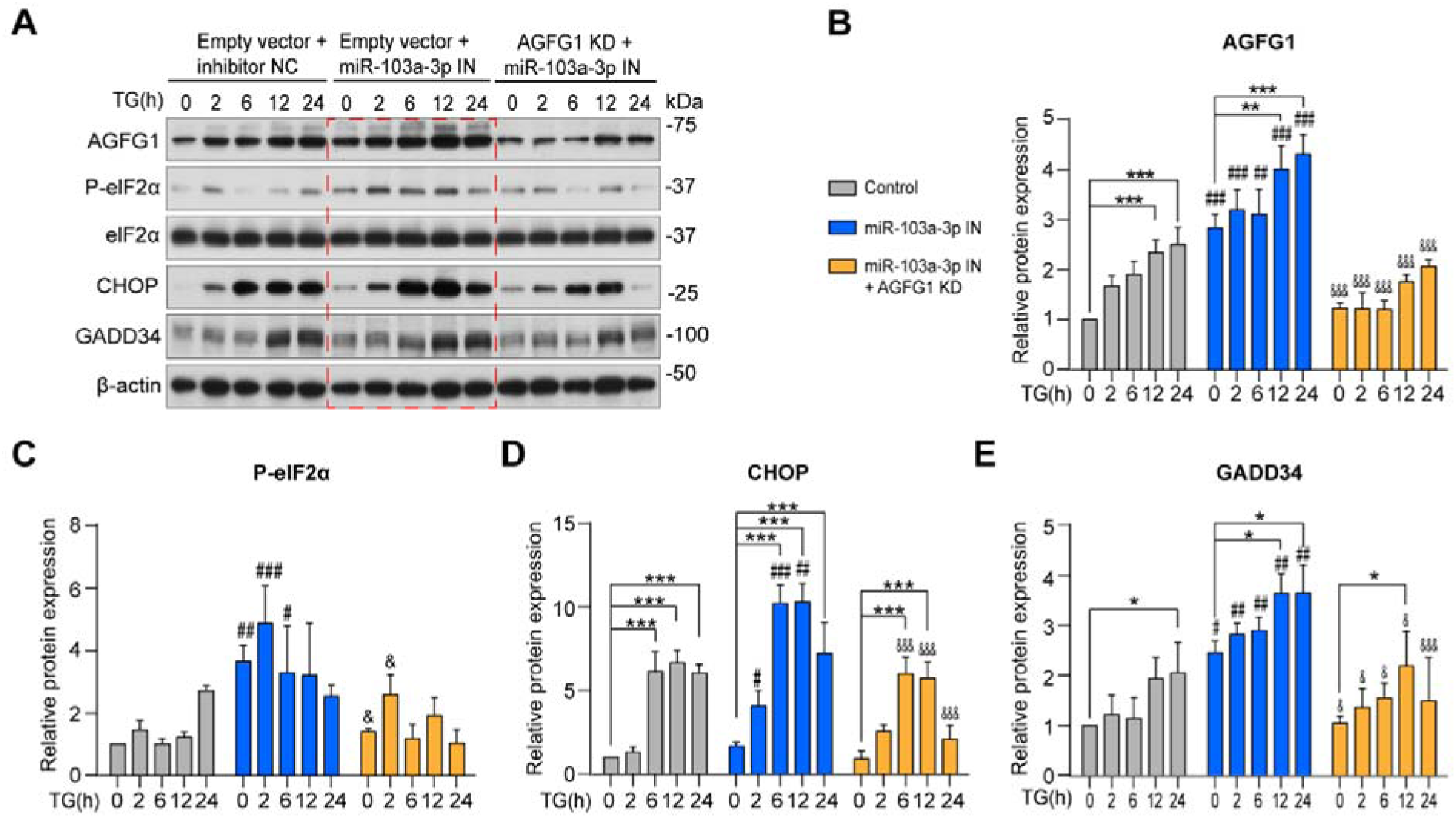
AGFG1 knockdown rescues ER stress induced by miR-103-3p inhibition. AGFG1 KD and vector control stable cells were transfected with miR-103a-3p inhibitor for 48 h, and then treated with 1 μM TG for 2, 6, 12 and 24 h. (**A**) Representative Western blot and quantification of (**B**) AGFG1, (**C**) P-eIF2α/eIF2α, (D) CHOP, and (**E**) GADD34. Data are presented as the mean ± SD (*n* = 3 per group). \**p* < 0.05, \*\**p* < 0.01, and \*\*\**p* < 0.001 relative to 0 h TG; ^#^*p* < 0.05, ^##^*p* < 0.01, and ^###^*p* < 0.001 relative to corresponding same timepoint of TG treatment in vector control transfected with inhibitor control; ^&^*p* < 0.05, and ^&&&^*p* < 0.001 relative to corresponding same timepoint of TG treatment in vector control transfected with miR-103a-3p inhibitor.

### AGFG1 localises to clathrin-positive endocytic vesicles in neuronal cells

AGFG1 has been implicated in clathrin-mediated endocytosis (CME), a process initiated by adaptor protein 2 (AP2) and clathrin assembly at the plasma membrane and Golgi apparatus^44–46^. To characterise the subcellular localisation of AGFG1, we performed immunofluorescence analysis in SH-SY5Y cells. Endogenous AGFG1 co-localised with clathrin light chain (CLC) and AP2 adaptor complex (**Fig. 5A-B**). Similar pattern of co-localization were observed in mouse primary cortical neurons and human induced pluripotent stem cell (iPSC)-derived neurons, indicating that endocytic localization of AGFG1 is conserved across neuronal systems (**Fig. 5C-D**). We further examined the intracellular distribution of GFP-tagged AGFG1 in SH-SY5Y cells. GFP-AGFG1 was predominantly localised to the perinuclear Golgi region and peripheral vesicle structures (**Fig. 5E**). Co-expression of GFP-AGFG1 with RFP-tagged AP2 or CLC revealed substantial co-localisation and co-trafficking of AGFG1 with clathrin-coated vesicles (**Fig. 5F and Suppl mov.S1**), supporting the association of AGFG1 with clathrin-mediated endocytic (CME) structures.

**Figure 5.**
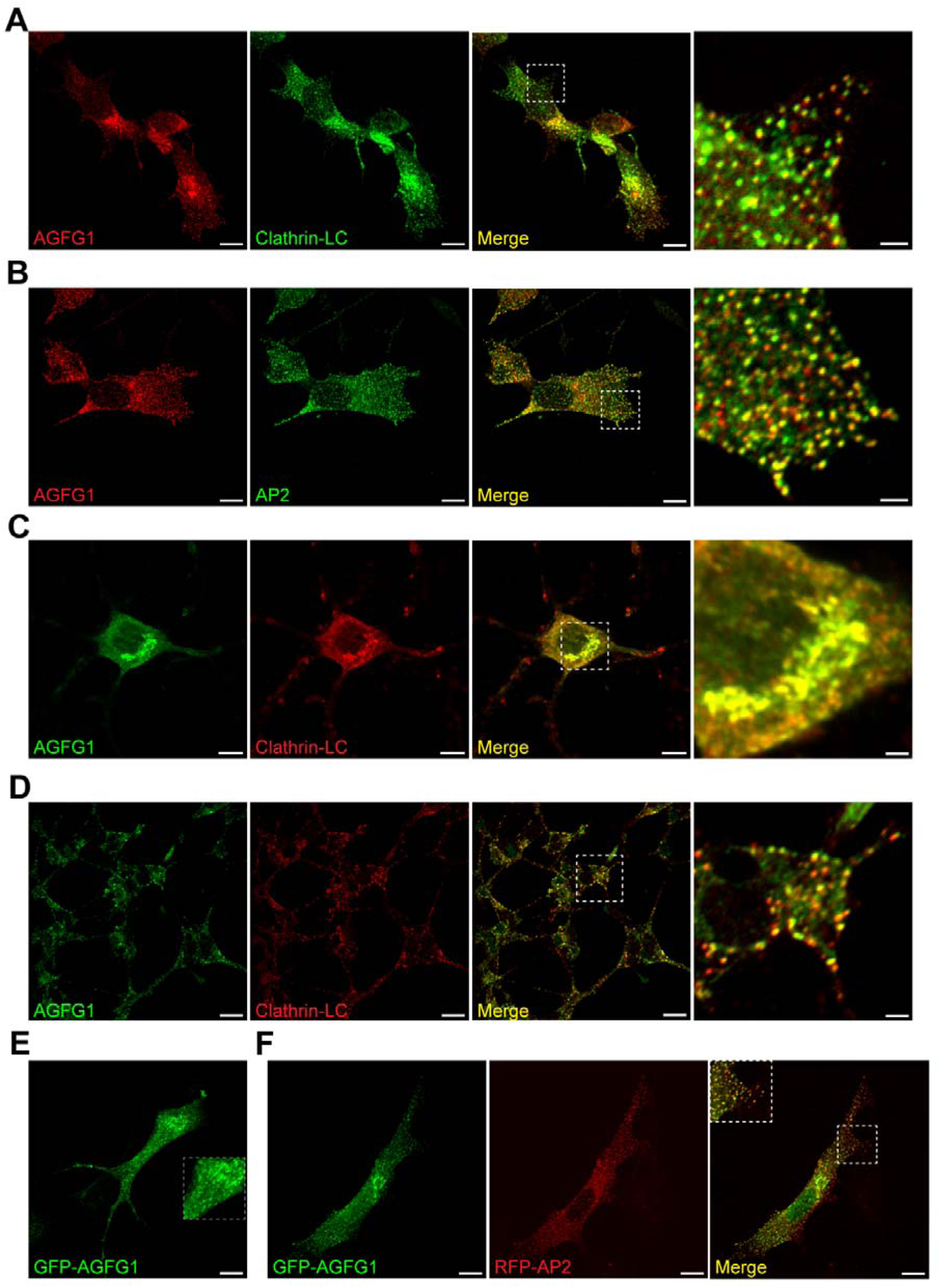
AGFG1 locolised to clathrin-positive endocytic vesicles in neuronal cells. Immunofluorescence analysis in SH-SY5Y neuroblastoma cells demonstrates co-localization of endogenous AGFG1 with (**A**) clathrin light chain (CLC) and (**B**) the adaptor AP2, key components of clathrin-mediated endocytosis (CME). Endogenous AGFG1 also co-localises with CLC in (**C**) mouse primary cortical neurons and (**D**) human iPSC-derived neurons, indicating this association is conserved across species and neuronal systems. (**E**) Overexpressed GFP-AGFG1 in SH-SY5Y cells localises at peri-nuclear Golgi and peripheral vesicle structures. (**F**) Co-expression of GFP-AGFG1 and RFP-AP2 in SH-SY5Y cells reveals their co-localization on clathrin-coated vesicles (CCVs). Scale bar: 10 µm (original views in **A-F**), 2 µm (enlarged view in **A-D**).

To further investigate the relationship between AGFG1 and clathrin-mediated membrane trafficking, we analysed primary cortical neurons derived from newborn of SJ-KI^RQ^ mice and wild-type (WT) littermate^47,48^. SJ-KI^RQ^ mice harbor the human early-onset Parkinsonism R258Q mutation (corresponding to R259Q in mouse Synaptojanin (SJ1) and several endocytic proteins involved in CME are abnormally accumulated at presynaptic terminals in KI neurons due to impaired clatrhin uncoating and synaptic vesicle recycling^47,48^. In WT neurons, endogenous AGFG1 co-localised with CLC and auxilin (**Supplementary Fig. S2**), a key uncoating factor required for clathrin-coated vesicle (CCV) disassembly, mainly at perinuclear Golgi region. In contrast, auxilin is re-located from soma to synapses and clustered at presynaptic terminals in SJ-KI^RQ^ neurons, suggesting impaired synaptic vesicle recycling. However, the somatic/Golgi distribution pattern of AGFG1 is not significantly changed (**Supplementary Fig. S2**). These findings suggest that AGFG1 may play a role in regulating clathrin-mediated Golgi and endocytic trafficking at soma instead of synaptic vesicle recycling at the terminals in primary neurons.

### ER stress disrupts AGFG1-associated endocytic dynamics in Parkinson’s disease models

To determine whether AGFG1-associated CME dynamics are altered by ER stress, we performed live-cell imaging of SH-SY5Y cells expressing AGFG1 and clathrin to track the dynamics of AGFG1-clathrin double-positive vesicles. TG treatment progressively reduced the formation and trafficking of AGFG1-clathrin double-positive vesicles, indicating impaired AGFG1-associated endocytic dynamics during ER stress (**Fig. 6A-B and Supplementary mov.S2**). We next examined CME-associated proteins^49^ in the SN of Parkinson’s disease. Protein levels of synaptojanin (SJ1), auxilin, and AP2-µ2 were significantly increased in Parkinson’s disease compared with controls, whereas CLC levels remained unchanged (**Fig. 6C-D**), suggesting potential compensatory upregulation within the endocytic pathway. Concurrently, tyrosine hydroxylase (TH) levels were significantly reduced, whereas-syn levels were significantly increased in SN of Parkinson’s disease (**Fig. 6E-F**), consistent with dopaminergic neurodegeneration and synuclein pathology in Parkinson’s disease.

**Figure 6.**
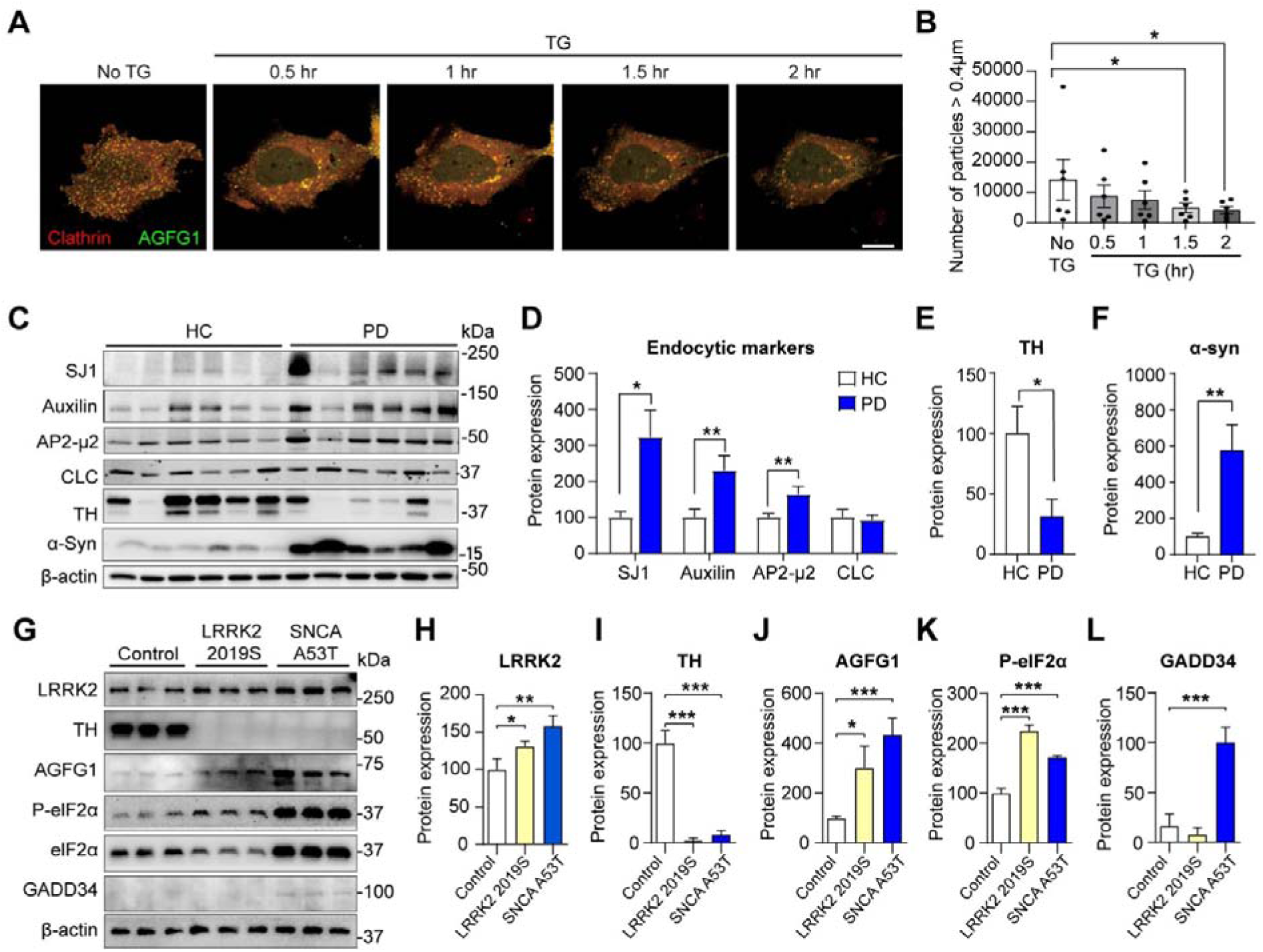
ER stress disrupts AGFG1-associated endocytocytic dynamics in Parkinson’s disease models. (**A-B**) Live-cell imaging of SH-SY5Y cells showed the dynamic trafficking of AGFG1-positive and clathrin-positive vesicles (clathrin-coated vesicles, CCVs). TG treatment progressively reduced the formation and trafficking of AGFG1-clathrin-positive vesicles. Scale bar: 20 µm (**C-F**) Representative Western blot and quantification of (**D**) CME-associated endocytic markers, (**E**) TH, and (**F**) □syn protein levels in the SN of Parkinson’s disease and HC (*n* = 6 per group). Parkinson’s disease showed increased protein levels of CME-associated endocytic protein including SJ1, Auxilin and AP2-µ2, accompanied by reduced TH levels and elevated-syn levels. (**G-M**) Neurons derived from SNCA A53T mutation hPSCs and LRRK2 G2019S iPSC exhibited neurodegeneration and ER stress. Representative Western blot and quantification of (**H**) LRRK2, (**I**) TH, (**J**) AGFG1, (**K**) P-eIF2α, (**I**) GADD34 protein expression. Data are presented as mean ± SD (*n* = 3 per group). \**p* < 0.05, \*\**p* < 0.01, and \*\*\**p* < 0.001. Two-group and multi-group comparisons were evaluated using Student’s *t*-test and one-or two-way ANOVA with Tukey’s post hoc test, respectively.

To further investigate the association between AGFG1 dysregulation and ER stress in genetic models of Parkinson’s disease, we examined hiPSC-derived pan-neuronal cultures habouring LRRK2 G2019S or SNCA A53T mutations. Both mutant neurons exhibited significantly increased LRRK2 expression and reduced TH expression compared with their shared isogenic control (**Fig. 6G-I**), suggesting that altered expression of PD-associated and dopaminergic neuronal markers. Consistent with the elevated AGFG1 protein expression observed in SN of Parkinson’s disease (**Fig. 1C**), AGFG1 expression were significantly increased in SNCA A53T neurons but not in LRRK2 G2019S neurons (**Fig. 6J**). Notably, P-eIF2 levels were significantly increased in both mutant neurons (**Fig. 6K**), consistent with enhanced ER stress-associated UPR. However, GADD34 expression exhibited distinct expression patterns, with significant upregulation in SNCA A53T neurons but undetectable in LRRK2 G2019S neurons (**Fig. 6L**). As GADD34 promotes eIF2 dephosphorylation through a negative feedback regulation mechanism^51,52^, its increaseed expression in SNCA A53T neurons may reflect activation of a compensatory response to sustained stress signaling, despite persistent elevation of P-eIF2. In contrast, elevated P-eIF2 in LRRK2 G2019S neurons without corresponding AGFG1 and GADD34 upregulation suggests that distinct regulatory mechanisms may contribute to stress responses in this genetic model. Together, these findings highlight a potential association between AGFG1 upregulation and ER stress-associated signaling in a context of mutant □synculein, suggesting mutation-dependent differences in cellular stress responses.

### ASO-mediated AGFG1 suppression attenuates ER stress and dopaminergic neurodegeneration in vivo

To evaluate the therapeutic potential of AGFG1 inhibition, we screened 50 antisense oligonucleotides (ASOs) targeting AGFG1 in SH-SY5Y cells. Among the candidates tested, ASO1 and ASO2 significantly reduced AGFG1 protein levels after 48 h of treatment, indicating efficient target knockdown (**Fig. 7A-B**). AGFG1-ASO treatment also significantly reduced TG-induced ER stress markers, including P-eIF2, CHOP and GADD34 (**Fig. 7C-E**). These findings were consistent with the effect observed following stable AGFG1 knockdown (**Fig. 3**), supporting a critical role for AGFG1in ER stress regulation.

**Figure 7.**
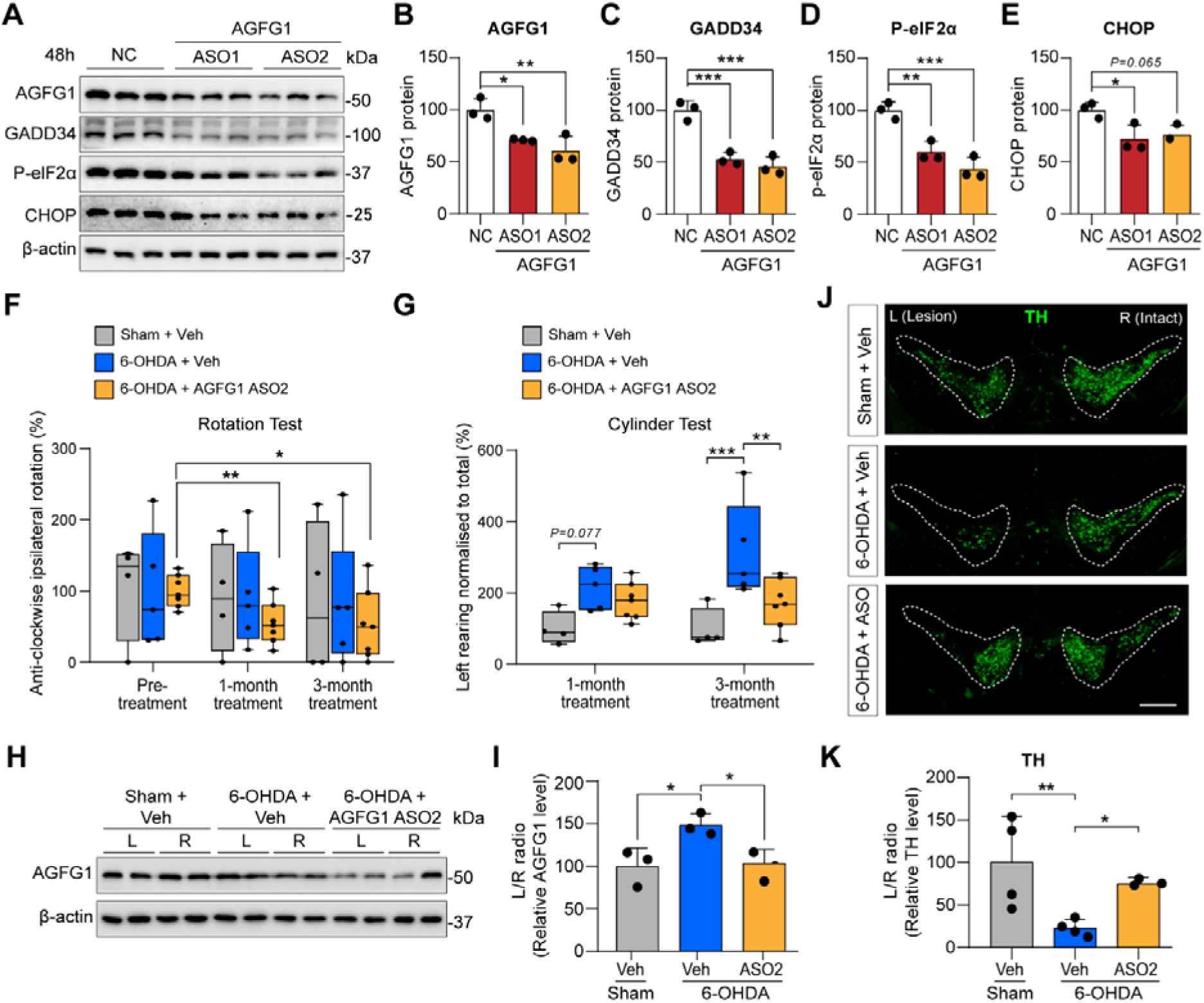
ASO-mediated AGFG1 suppression attenuates ER stress signaling and dopaminergic neurodegeneration in 6-OHDA-induced Parkinson’s disease mouse model. (A-E) SH-SY5Y cells were transfected with 100 nM of AGFG1-ASO1,-ASO2, and control for 48 h. **(A)** Representative Western blot and quantification of protein levels of (B) AGFG1, **(C)** GADD34, **(D)** P-eIF2α, and **(E)** CHOP. Data are mean ± SD (*n* = 3 per group). \**p* < 0.05, \*\**p* < 0.01, and \*\*\**p* <0.001 by one-way ANOVA with Tukey’s post hoc test. (**F-G**) *In vivo* validation of 6-OHDA Parkinson’s disease mouse model. 6-OHDA (4 μg) was stereotaxically injected into left striatum of two-month-old C57 (day 0). AGFG1-ASO2 (2 µg) was then administered into the left SN on day eight. Motor behavior was assessed using (**F**) the rotation test and (**G**) the cylinder test at one-month and three-month post-treatment. Data are mean ± SD (vehicle, *n* = 4; ASO-treated, *n* = 7). (**H-I**) Representative Western blotting and quantification of AGFG1 expression in left and right striata, expressed as L/R ratios. (**J-K**) Immunofluorescence staining for TH in the substantia nigra pars compacta (SNpc; scale bars, 500 μm), with quantification of total TH-positive cells expressed as L/R ratios. Data are means ± SD (*n* = 6 per group). \**p* < 0.05, \*\**p* < 0.01 by one-way ANOVA with Tukey’s post hoc test.

We next investigated the therapeutic potential of AGFG1-ASO2 in a unilateral 6-hydroxydopamine (6-OHDA) mouse model of Parkinson’s disease. Based on its greater knockdown efficiency, AGFG1-ASO2 was stereotaxically administered into the SN of 6-OHDA-lesioned mice. In the amphetamine-induced rotation test, vehicle-treated 6-OHDA mice exhibited pronounced ipsilateral rotations, indicating severe dopaminergic damage (**Fig. 7F**). AGFG1-ASO2 treatment significantly reduced ipsilateral rotations at both one and three months post-treatment, indicating sustained improvement in dopaminergic motor function (**Fig. 7F**). Similarly, 6-OHDA lesioning significantly impaired contralateral forelimb use in the cylinder test, whereas AGFG1-ASO2 treatment significantly improved the forelimb use, with a greater recovery observed at three months (**Fig. 7G**).

At the molecular level, 6-OHDA lesioning induced a significant upregulation of AGFG1 protein expression in the SN, whereas AGFG1-ASO2 treatment significantly suppressed this increase (**Fig. 7H-I**). Immunohistochemical analysis further demonstrated a marked reduction in TH-positive neurons in the SN of 6-OHDA lesioned-mice, which was significantly attenuated by AGFG1-ASO2 treatment (**Fig. 7J**). Consistent with these findings, the ratio of TH levels in the lesioned (left) SN relative to the intact (right) SN was significantly reduced in vehicle-treated 6-OHDA mice, whereas AGFG1-ASO2 treatment significantly increased TH levels in the lesioned side, indicating neuroprotective efficacy.

Collectively, these findings demonstrate that ASO-mediated suppression of AGFG1 attenuates ER stress *in vitro* and reduces dopaminergic neurodegeneration and motor deficits in the 6-OHDA mouse mouse models. These findings provide initial proof-of-concept for the neuroprotective potential of AGFG1 inhibition.

## Discussion

Dysregulated miRNAs expression has been widely reported in Parkinson’s disease across patients samples, animal models and cellular systems^16^. However, miRNAs profiles are highly context-dependent and may vary according to disease stage, tissue source, cellular composition and treatment status. In this study, miR-103a-3p was significantly reduced in both plasma and SN of Parkinson’s disease (**Fig. 1A-B**), supporting an association between miR-103a-3p dysregulation and Parkinson’s disease pathology^53–55^. Previous studies have reported increased miR-103a-3p expression in mitophagy-induced Parkinson’s disease models^56^ and in peripheral blood mononuclear cells from L-dopa-treated patients with Parkinson’s disease^57^, whereas no significantly difference was observed in drug-naïve patients^57^. These apparently divergent findings may reflect differences in experimental models, sample types, disease stage, and treatment exposure, highlighting the importance of biological and clinical context when interpreting miRNA expression across studies. Importantly, our findings of reduced miR-103a-3p in both peripheral plasma and affected brain tissue suggests that altered miR-103a-3p levels is associated with PD, although larger and clinically stratified cohorts will be required to determine its relationship with disease stage and treatment status.

Mechanistically, we identidied AGFG1 as a direct target and downstream effector of miR-103a-3p. (**Fig. 1D-E**). Reduced miR-103a-3p expression in SN of Parkinson’s disease SN was accompanied with increased AGFG1 protein levels without a corresponding change in AGFG1 mRNA (**Fig. 1B-C and Supplementary Fig. S1B**). A similar inverse relationship was observed in SH-SY5Y cells following miR-103a-3p inhibition, which increased AGFG1 protein without significantly altering its mRNA level (**Fig. 1H-I and Supplementary Fig. S1E**). Together with the luciferase reporter assay (**Fig. 1E**), these findings support post-transcriptional regulation of AGFG1 by miR-103a-3p. Increased AGFG1 expression has also been reported in a large human SN transcriptomic dataset^58^, although further experimental validation across independent cohorts is warranted. Previous studies have identified several targets through which miR-103a-3p may regulate proteolysis, autophagy and cell survival, including Parkin, phosphatase and tensin homolog (PTEN), autophagy related 5 (Atg5), and high mobility group box 1 (HMGB1)^59–62^. Our data extend these observations by identifying AGFG1 as a previously unrecognized target through which miR-103a-3p may intersect with endocytic trafficking and ER stress pathways. Notably, although AGFG1 has previously been predicted *in silico* as a potential miR-103a-3p target in PD^57^, its functional involvement in neurodegenerative processes has not been established.

AGFG1 is an endocytic protein predominantly localised to the perinuclear region, plasma membrane and Golgi apparatus, and has been implicated in the CME^44–46^. In this study, AGFG1 co-localised with clathrin and the AP2 adaptor complex in SH-SY5Y cells, primary cortical neurons, and human iPSC-derived neurons (**Fig. 5**), supporting an association between AGFG1 and clathrin-coated structures across neuronal systems. Live-cell imaging further revealed a progressive reduction in AGFG1-clathrin double-positive vesicles formation following induction of ER stress by TG (**Fig. 6A-B**), indicating that sustained ER stress is accompanied by impaired AGFG1-associated endocytic dynamics by disrupting the CCV formation or maturation. Given that impaired CME can prolong the plasma membrane residence of CME cargo proteins^63^, potentially altering the transport activity and receptor signaling. Consistent with this possibility, we observed increased expressions of AGFG1 (**Fig. 1C**), AP2-µ2, SJ1, and auxilin in the SN of Parkinson’s disease compared with HC (**Fig. 6C-D**). The coordinated increased in these endocytic proteins may reflect an adaptive response to altered endocytic trafficking, with increased AGFG1 and AP2-µ2 supporting vesicle assembly and elevated uncoating factor SJ1 and auxilin facilitating the clearance of non-productive clathrin assemblies. However, whether these changes are compensatory or contribute directly ro endocytic dysfunction remains to be determined. Collectively, these findings support an association between AGFG1 dysregulation, altered CME machinery, and PD pathology, while further studies directly measuring endocytic flux will be required to define the functional consequences of AGFG1 regulation.

ER stress and dysregulation of UPR have been implicated in dopaminergic neurodegeneration in Parkinson’s disease^64^. Our findings identify the miR-103a-3p/AGFG1 axis as a potential regulator of ER stress responses. Overexpression of miR-103a-3p attenuated TG-induced P-eIF2α, and downstream induction of CHOP and GADD34^11,65–67^ (**Fig. 2A-D**), whereas miR-103a-3p inhibition exacerbated ER stress and promoted pro-apototic signaling^61^ (**Fig. 2E-H**). Importantly, AGFG1 knockdown attenuated the enhanced ER stress signaling induced by miR-103a-3p (**Fig 4**), supporting AGFG1 as a functional downstream mediator of miR-103a-3p-dependednt ER stress regulation. Consistently, both genetic knockdown (**Fig. 3G-K**) and ASO-mediated suppression of AGFG1 (**Fig. 7A-E**) attenuated TG-induced ER stress responses. Together, these findings further supporting the involvement of AGFG1 in UPR regulation.

The relevance of these findings to genetic Parkinson’s disease was further supported by distinct AGFG1 and ER stress-associated expression patterns in SNCA A53T and LRRK2 G2019S neurons. Although both models exhibited significantly increased expression of P-eIF2α, AGFG1 and GADD34 upregulation was observed only in SNCA A53T neurons (**Fig. 6J-L**). Previous studies reported that mutant α-synuclein discrupts ER-associated protein homeostasis and activates ER stress responses in human iPSC-derived neurons^68,69^, whereas LRRK2 G2019S has been associated with altered ER calcium homeostasis and RAB8A-dependent endolysosomal trafficking^70,71^. These distinct pathogenic processes may contribute to the differential stress-response profiles observed between these two genetic models. In particular, the selective upregulation of AGFG1 in SNCA A53T neurons raise the possibility that the AGFG1 dysregulation may be more closely associated with α-synuclein-related cellular stress. Notably, GADD34 promotes eIF2α dephosphorylation of through a stress-induced negative feedback mechanism that facilitates recovery of protein synthesis^51,72^. The concomitant increased P-eIF2α and GADD34 expression in SNCA A53T neurons may therefore reflect engagement of this feedback response during persistent ER stress-associated signaling. In contrast, LRRK2 G2019S neurons exhibited increased P-eIF2α without corresponding AGFG1 or GADD34 upregulation (**Fig. 6J-L**), suggests that additional mechanisms may contribute to stress signaling in this mdoel. Together, these findings highlight mutant-dependent differences in AGFG1 expression and ER stress-associated signaling, supporting further investigation of AGFG1 in □synuclein-associated pathology and its potential contribution to UPR regulation.

One potential mechanism linking AGFG1-dependent trafficking to ER homeostatis may involve vesicle-associated membraine protein 7 (VAMP7). AGFG1 has been reported to interact with the longin domain of VAMP7 during the early stage of CME and to contribute to the assembly and fusion of CCV at the plasma membrane^32,45^. Silencing of AGFG1 has also been reported to increase VAMP7 accumulation at the plasma membrane^73^, indicating that AGFG1 contributes to VAMP7 trafficking and recycling. Conversely, VAMP7 knockdown has been associated with reduced CCV fusion, impaired secretion of ER-associated components and ER stress^74^. Based on these observations, sustained AGFG1 upregulation in Parkinson’s disease could potentially perturb VAMP7-dependent membrane trafficking and thereby influence ER homeostasis and stress signaling. However, as VAMP7-dependent trafficking was not directly examined in this study, determining whether VAMP7 or other CME components mediate the effects of AGFG1 on ER stress will thererefore required for future investigation.

The therapeuticic relevance of AGFG1 was further supported by both genetic and pharmacological approaches. Suppression of AGFG1 reduced ER stress in cellular models, while stereotaxic administration of AGFG1-targeting ASO reduced motor deficits and, preserved nigral dopaminergic neurons in the 6-OHDA mouse model (**Fig. 7**). These findings provide proof-of-concept that reducing AGFG1 expression can protect the nigrostriatal dopaminergic system against neurodegenerative injury. Together with the dysregulation of the miR-103a-3p/AGFG1 axis observed in human Parkinson’s disease tissue and neuronal models, these results support AGFG1 as a potential therapeutic target for mitigating ER stress-associated dopaminergic neurodegeneration. Nevertheless, the 6-OHDA model induces relatively rapid toxin-mediated dopaminergic injury and does not recapitulate the progressive, cell-autonomous dysfunction and-syn-associated pathology of the human disease^75,76^. Thus, while our findings estabalish the neuroprotective potential of AGFG1 suppression *in vivo*, future studies using complementary models that incorporate progressive-syn pathology will be important to determine the broader therapeutic applicability of targeting AGFG1 in Parkinson’s disease.

In summary, our findings identify a previously unrecognised miR-103a-3p/AGFG1 regulatory axis linking altered endocytic trafficking with ER stress in Parkinson’s disease. Reduced miR-103a-3p expression is associated with increased AGFG1 protein expression, while manipulation of the miR-103a-3p/AGFG1 axis modulates ER stress responses. Pharmacological suppression of AGFG1 further attenuated ER stress and mitigated dopaminergic neurodegeneration and motor deficits in 6-OHDA mouse model, providing proof-of-concept for its neuroprotective potential. Together these findingss provide new insight into the intersection between miRNA dysregulation, endocytic trafficking, and ER stress in Parkinson’s disease and support further investigation of AGFG1 as a potential therapeutic target for Parkinson’s disease.

## Availability of data and materials

Data that supports the findings of this study will be available upon request.

## Acknowledgements

We would like to thank all our patients and their families, the staff of all clinics and hospital for their valuable contributions to the study.

This study was supported by Singapore Ministry of Health’s National Medical Research Council through Clinician Scientist-Individual Research Grant (CIRG21nov-0001, CIR23jul-0006), Open Fund Investigator Research Grant (OFIRG23jul-0075), and Open Fund-Large Collaborative Grant (OFLCG24may-0026). Additional support was provided by the SingHealth Duke-NUS Academic Medicine Research Grant (AM/TP114/2026 (SRDUKAMR2614)).

## Competing interests

Authors Li Zeng, Haitao Tu, and Zhi Wei Zhang declare that a patent application (#10202503222V) has been filed for the therapeutic targeting of AGFG1 using antisense oligonucleuotides (ASOs) identified in this study. All other authors declare no conflicts of interest.

